## Supplemental Information for "AIRRscape: an interactive tool for exploring B-cell receptor repertoires and antibody responses"

**S9 Fig. Dengue convergent clonotypes to Parameswaran (2018) motif ARQIGNWFDP similar to CF1 (Zanini 2018).** An 80% identity threshold is used to calculate convergence. Tips are colored by dataset source. Purple tips are plasmablast sequences reported by Zanini (2018) isolated from two Colombian patients (d13 and d20), blue tips are antibody sequences from the BCR repertoire of patient d13, and gold tips are antibody sequences from a cohort of Nicaraguan patient BCR repertoires.

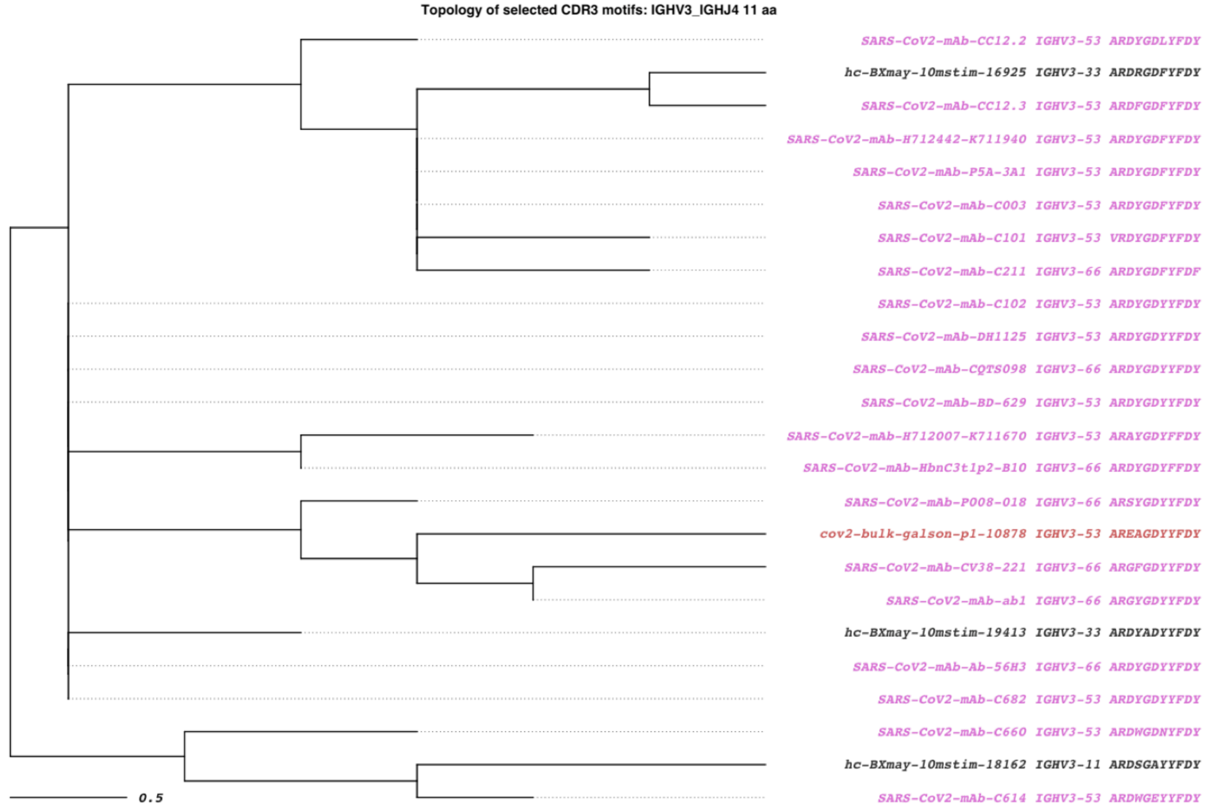


**S1 Fig. SARS-CoV-2 convergent clonotypes to mAb C102 in the 3_4_11 bin.** An 80% identity threshold is used to calculate convergence. Tips are colored by dataset source. Purple tips are published anti-COVID-19 antibodies from 12 different studies, dark gray tips are antibody sequences from a healthy donor BCR repertoire, and orange through brown shaded tips are antibody sequences from COVID-19 patient BCR repertoires.

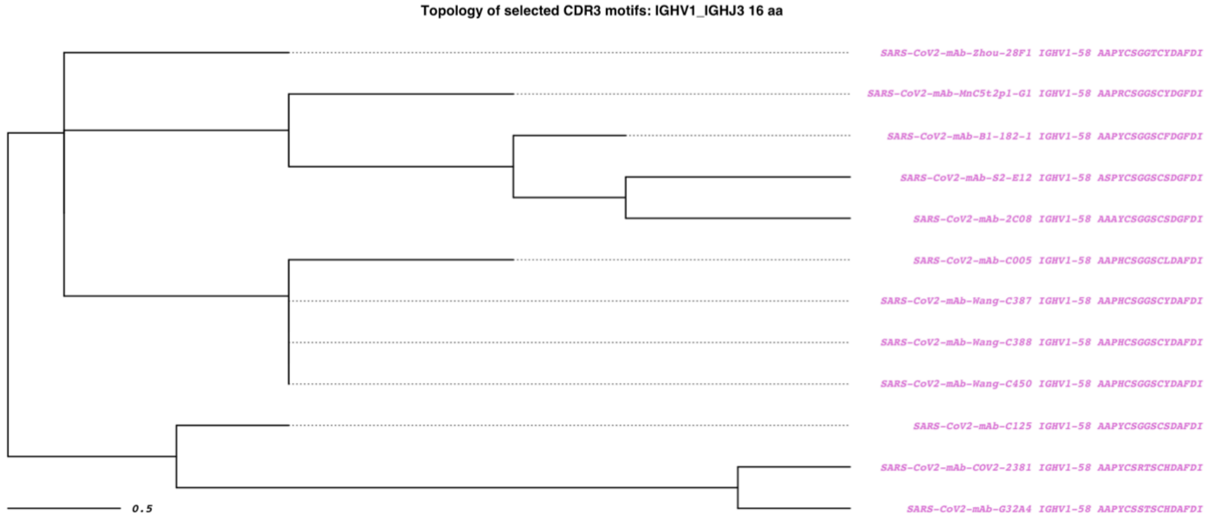


**S2 Fig. SARS-CoV-2 convergent clonotypes to mAb C125 in the 1_3_16 bin.** An 80% identity threshold is used to calculate convergence. Tips are colored by dataset source. Purple tips are published anti-COVID-19 antibodies from 7 different studies, dark gray tips are antibody sequences from a healthy donor BCR repertoire, and orange through brown shaded tips are antibody sequences from COVID-19 patient BCR repertoires.

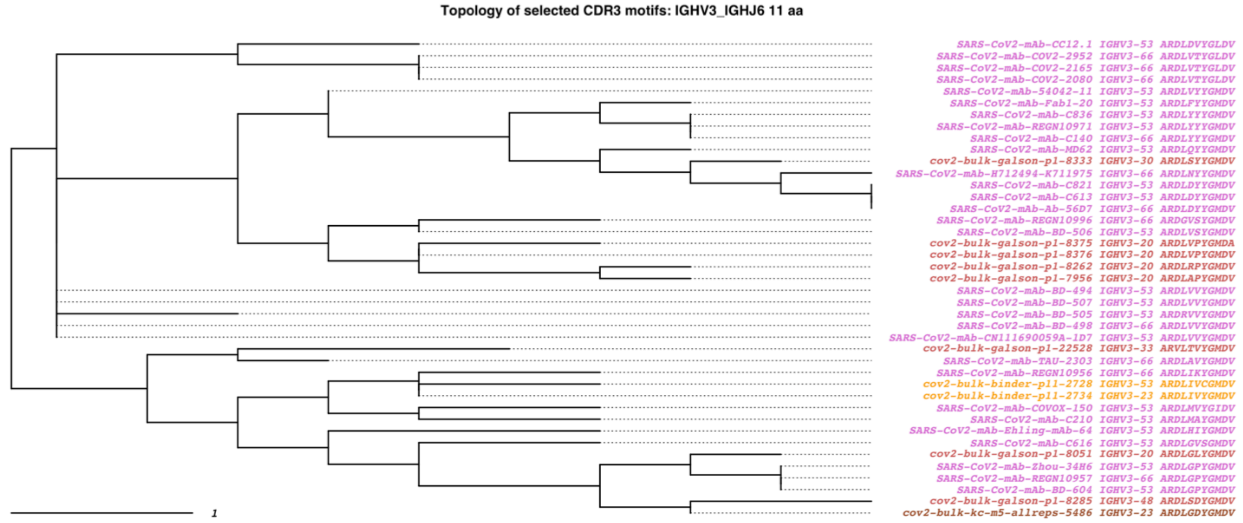


**S3 Fig. SARS-CoV-2 convergent clonotypes to mAb BD-498 in the 3_6_11 bin.** An 80% identity threshold is used to calculate convergence. Tips are colored by dataset source. Purple tips are published anti-COVID-19 antibodies from 15 different studies, dark gray tips are antibody sequences from a healthy donor BCR repertoire, and orange through brown shaded tips are antibody sequences from COVID-19 patient BCR repertoires.

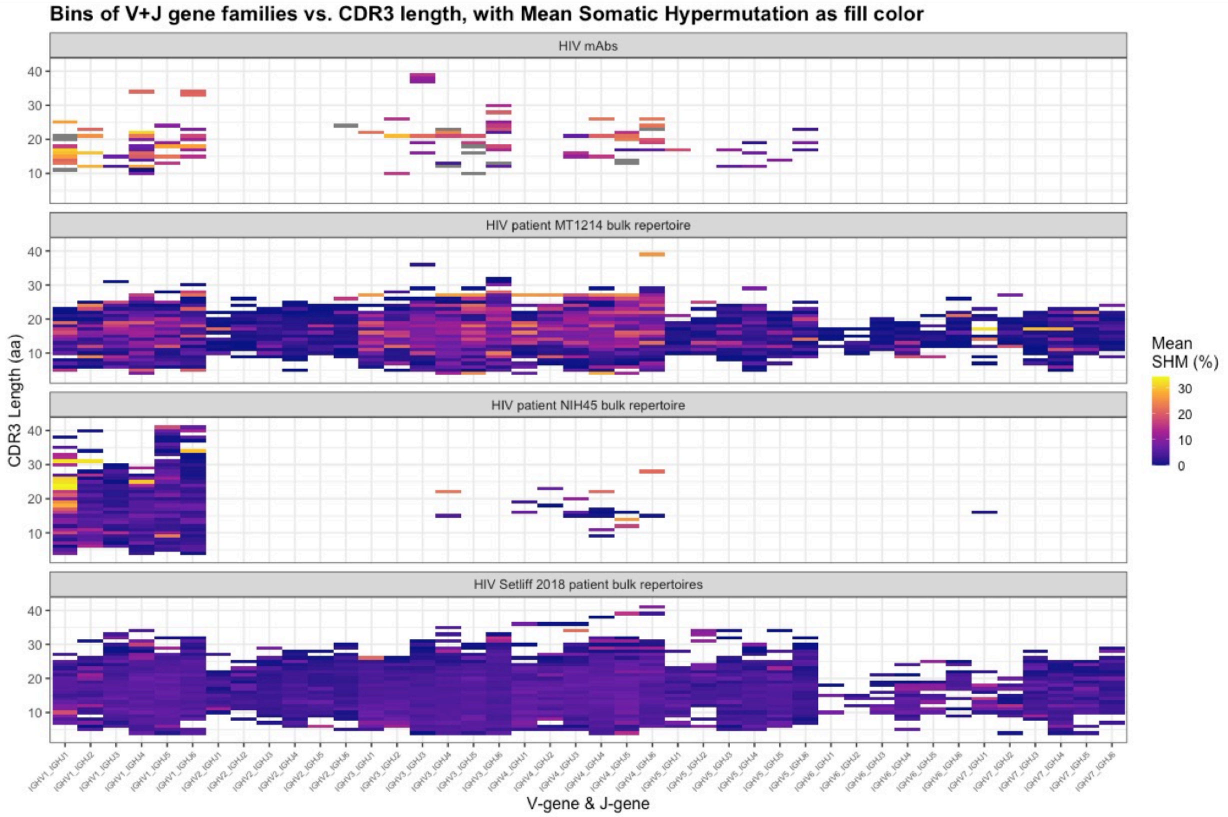


**S4 Fig. AIRRscape heatmaps comparing anti-HIV-1 antibodies and bulk BCR repertoires of eight HIV-1 patients.**

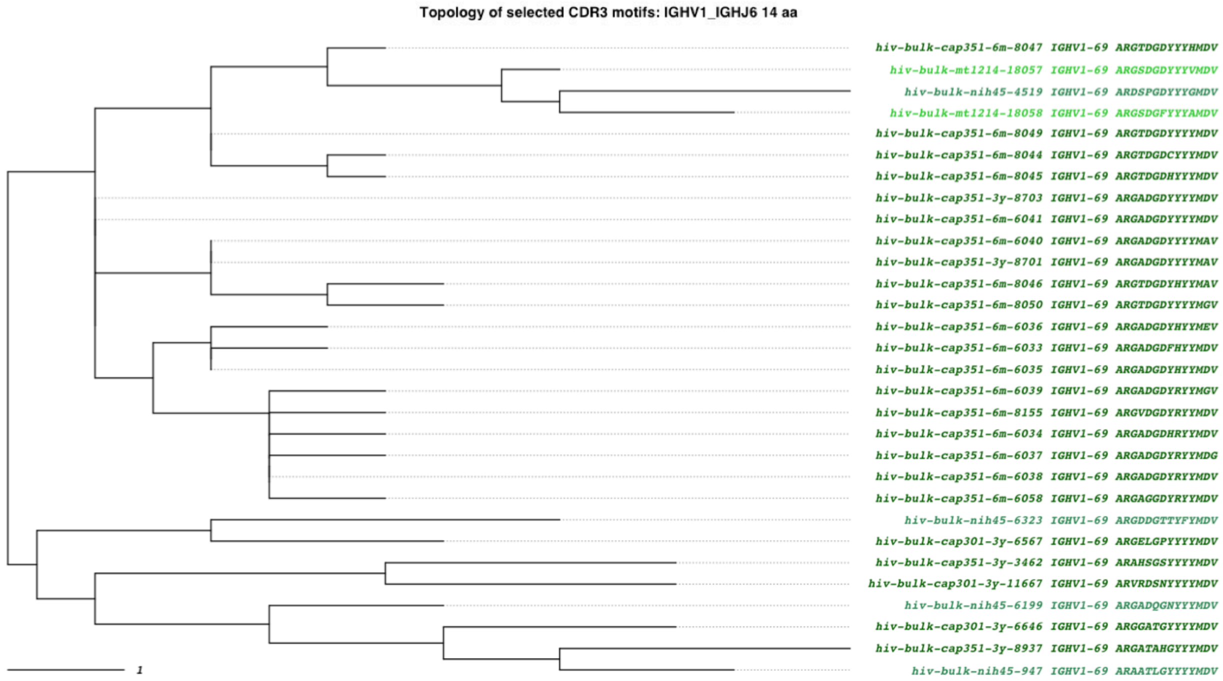


**S5 Fig. HIV-1 convergent clonotypes to mAb CAP351_6m_6041 (Setliff 2018; Fig 3) in the 1_6_14 bin.** A 70% identity threshold is used to calculate convergence. Tips are colored by dataset source. Green shaded tips are antibody sequences from HIV-1 patient BCR repertoires.

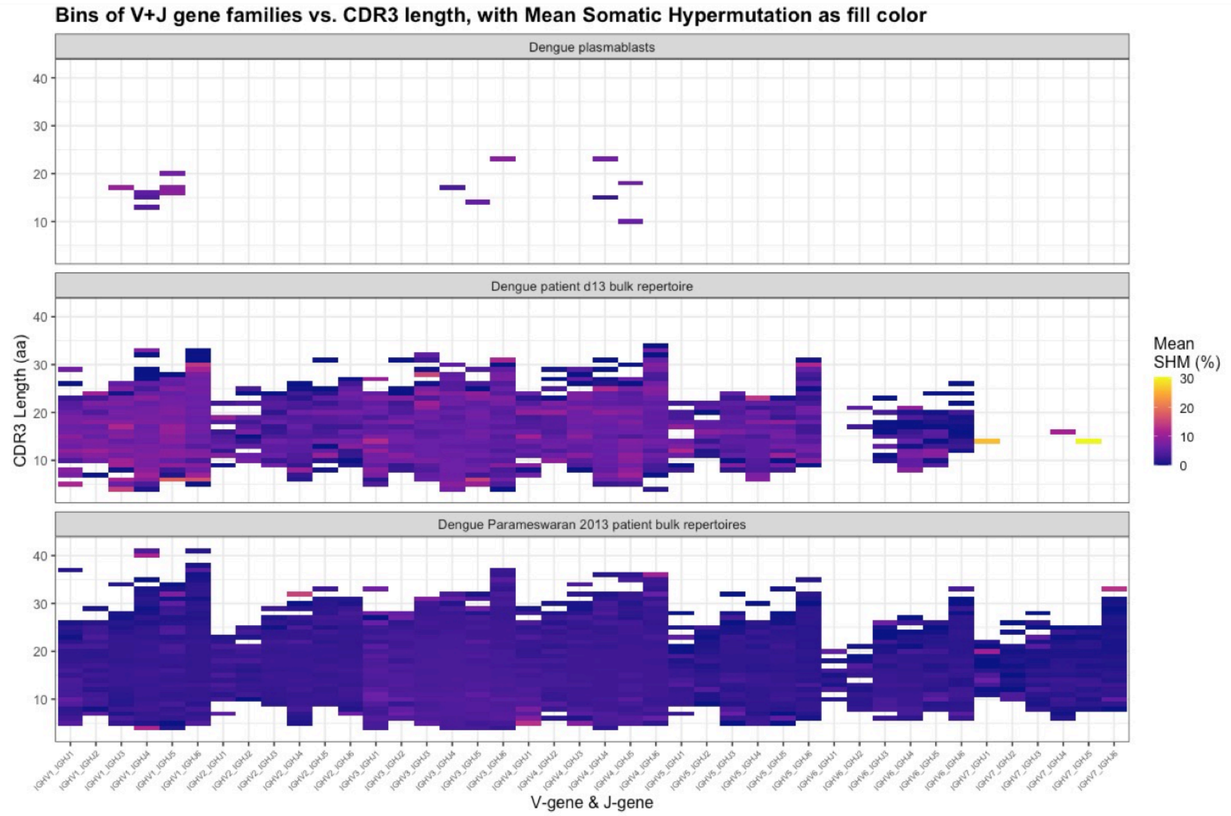


**S6 Fig. AIRRscape heatmaps comparing isolated dengue plasmablasts and bulk BCR repertoires of dengue patients from Colombian and Nicaraguan cohorts.**

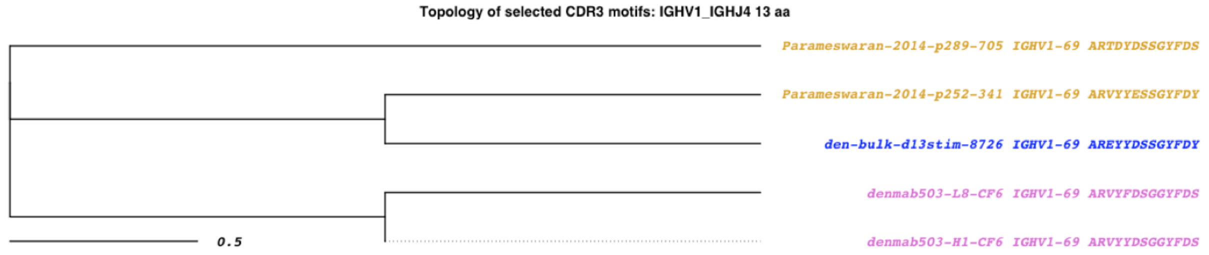


**S7 Fig. Dengue convergent clonotypes to CF6 (Zanini 2018).** An 80% identity threshold is used to calculate convergence. Tips are colored by dataset source. Purple tips are plasmablast sequences reported by Zanini (2018) isolated from two Colombian patients (d13 and d20), blue tips are antibody sequences from the BCR repertoire of patient d13, and gold tips are antibody sequences from a cohort of Nicaraguan patient BCR repertoires.

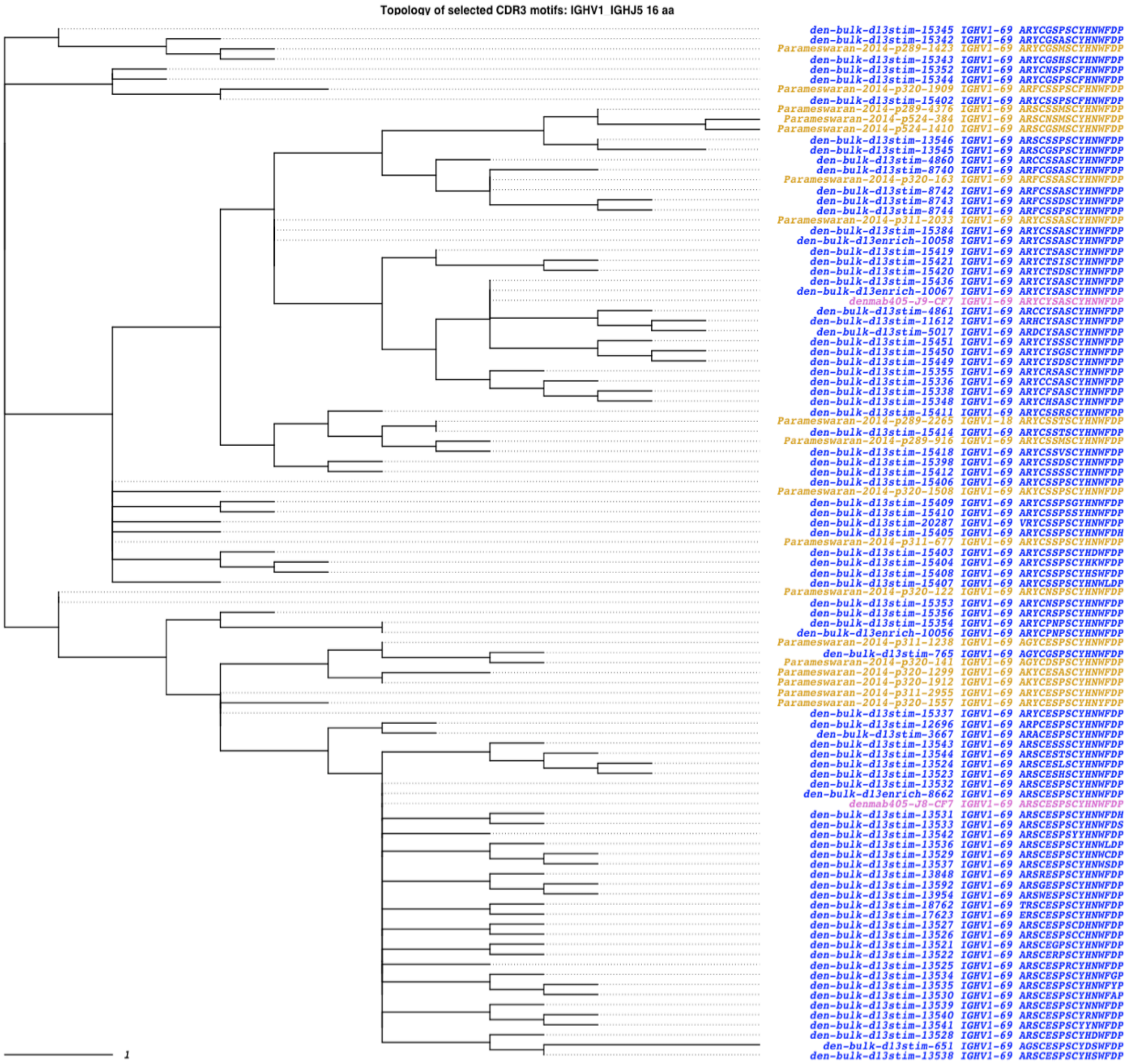


**S8 Fig. Dengue convergent clonotypes to CF7 (Zanini 2018).** An 80% identity threshold is used to calculate convergence. Tips are colored by dataset source. Purple tips are plasmablast sequences reported by Zanini (2018) isolated from two Colombian patients (d13 and d20), blue tips are antibody sequences from the BCR repertoire of patient d13, and gold tips are antibody sequences from a cohort of Nicaraguan patient BCR repertoires.

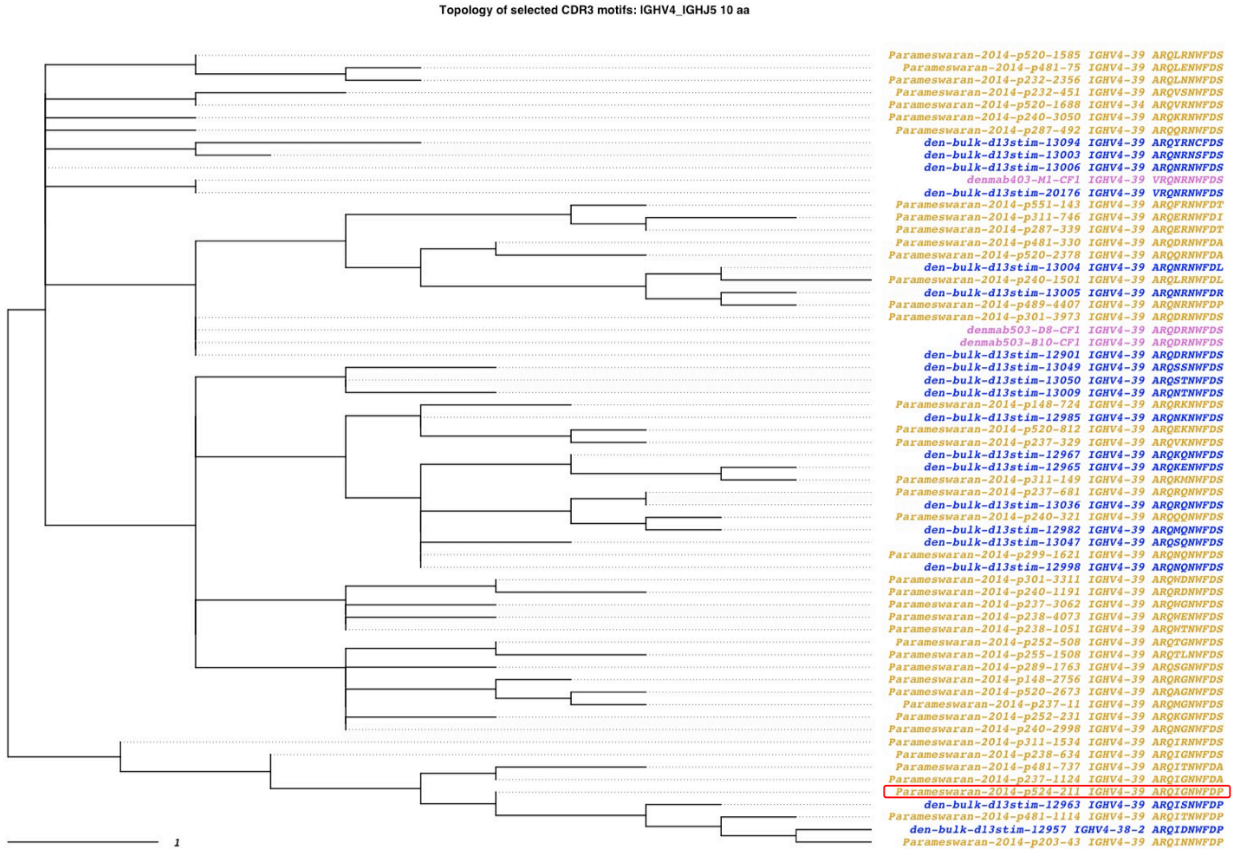


**S9 Fig. Dengue convergent clonotypes to Parameswaran (2018) motif ARQIGNWFDP similar to CF1 (Zanini 2018).** An 80% identity threshold is used to calculate convergence. Tips are colored by dataset source. Purple tips are plasmablast sequences reported by Zanini (2018) isolated from two Colombian patients (d13 and d20), blue tips are antibody sequences from the BCR repertoire of patient d13, and gold tips are antibody sequences from a cohort of Nicaraguan patient BCR repertoires.
